# Increased tau expression in the *APOE4* blood-brain barrier model is associated with reduced anti-tau therapeutic antibody delivery *in vitro*

**DOI:** 10.1101/2023.10.24.563706

**Authors:** Joanna M. Wasielewska, Rinie Bajracharya, Rebecca L. Johnston, Juliana C.S. Chaves, Alice Pébay, Lotta E. Oikari, Jürgen Götz, Rebecca M. Nisbet, Anthony R. White

## Abstract

Tau protein is a critical driver of neurodegeneration and an important drug target in Alzheimer’s disease (AD). Tau-specific immunotherapy has emerged as a promising treatment strategy for AD, however the therapeutic efficacy of anti-tau antibodies may be limited by their insufficient delivery across the blood-brain barrier (BBB). The apolipoprotein E4 allele (*APOE4*) is the strongest genetic risk factor for sporadic AD and is known to influence tau-mediated neurodegeneration. Interestingly, both tau and *APOE4* have been implicated in the cerebrovascular pathology observed in AD. Yet, the crosstalk between *APOE4* and tau at the level of the BBB and its consequences for anti-tau immunotherapeutics delivery, remain poorly understood. Here, we utilised *APOE3*- and *APOE4*-carrying human iPSC-derived induced brain endothelial-like cells (iBECs) as a sporadic AD BBB model, determined the levels of endogenous tau in iBECs, and explored the transport of two novel monoclonal anti-tau antibodies, RNF5 and RN2N, across the *in vitro* barrier. Our results demonstrate that *MAPT* gene transcription, tau protein levels and tau phosphorylation are increased in iBECs in an *APOE4*-related manner and are associated with reduced iBEC monolayer integrity and increased permeability to biologically inert fluorescent tracers. Additionally, elevated levels of intracellular tau in *APOE4* cells were accompanied by the reduced passive permeability of therapeutic anti-tau antibodies through the *APOE4* iBEC monolayer, which could be improved by the application of focused ultrasound and microbubble drug-delivery technology. Together, our study illustrates a new role for *APOE4* and tau in human iBECs with potential implications for BBB dysfunction and anti-tau therapeutic antibody delivery.

## INTRODUCTION

Alzheimer’s disease (AD) is a tauopathy characterised by the abnormal phosphorylation and accumulation of tau protein in the brain^1^. Currently, several anti-tau immunotherapies are being explored in clinical trials targeting AD^2–4^. Their clinical effectiveness, however, may be limited by the restrictive nature of the blood-brain barrier (BBB), which only allows an estimated 0.1% of therapeutic antibodies to reach the brain parenchyma^5–7^. Cerebrovascular brain endothelial cells (BECs) are the major cellular constituent of the BBB and are the first brain cells that peripherally administered immunotherapies encounter. Yet, little is known about their interaction with tau-specific therapeutics.

The BBB is altered in AD^8,9^, with a potential contribution of tau to cerebrovascular dysfunction^10–14^. Correspondingly, tau oligomers were found to accumulate in the cerebral microvasculature of AD patients and amyloid-depositing Tg2576 transgenic mice in close association with BECs^15^. Furthermore, microvascular abnormalities including spiralling morphologies and altered vessel diameter and density were identified in mouse models of tauopathy^16^. Similarly, tau-dependent cerebrovascular remodelling has been found at early Braak stages in human brain samples^17^. Tau was found to impact neurovascular coupling prior to the development of mature tau pathology and cognitive impairment in tau transgenic rTg4510 and PS19 mice^18^. In addition, strong relationships were reported between vascular tau and cognitive dysfunction in AD patients and with aging^19,20^. Studies in rTg4510 mice have revealed an association of perivascular tau with hippocampal BBB dysfunction, the latter’s integrity being recovered when tau levels were reduced, thereby establishing a causal link between BBB impairment and brain tau^21^. Recently, a study demonstrated that extracellularly transmitted tau oligomers of neuronal origin are internalised and accumulate in BECs contributing to the development of AD-like cerebrovascular dysfunction in mouse models of tauopathy^14^. Together, these studies suggest that pathogenic tau mediates BBB impairment possibly leading to complex drug-BBB interactions in AD. However, the role of endogenous tau within BECs has not yet been explored within that context.

The apolipoprotein E4 allele (*APOE4*) is the strongest genetic risk factor for sporadic AD, markedly accelerating tau pathology in brain parenchymal cells^22–25^. *APOE4* is also associated with increased BBB leakage and dysfunction^26–30^ and increased deposition of amyloid in cerebral vessel walls in the human brain and animal models^31–33^, as well as *in vitro* BBB systems^34^; however, its effect on tau accumulation at the BBB is incompletely understood. Additionally, while BBB dysfunction in AD is well established and generally (and possibly incorrectly) perceived as increased leakiness^8,9,29,35,36^, interestingly, increased brain uptake of peripherally administered therapeutics has not been reported^36–38^. Finally, the influence of the patients’ genetic profiles including the *APOE* genotype on the efficacy of tau immunotherapy delivery at the BBB is currently unknown.

Here, we identify increased levels of endogenous tau and tau phosphorylation in human *APOE4*-carrying induced BEC-like cells (iBECs) and assess their impact on anti-tau therapeutic antibody delivery *in vitro*. We show that the passive permeability of the novel anti-tau therapeutic antibodies RNF5 and RN2N is reduced at the barrier formed by *APOE4* cells as compared to *APOE3* controls, while this can be partially reversed by applying focused ultrasound and microbubble (FUS^+MB^) drug-delivery technology^39^. Furthermore, we demonstrate the presence of the investigated anti-tau antibodies in the *APOE4* iBEC monolayer, suggesting that the potential interaction of the antibodies with endogenous tau in BECs may be limiting antibody permeability at the BBB. While these findings should be complemented by further investigation *in vivo*, our study highlights a critical step towards demonstrating a role for the cross-talk of *APOE4* and tau at the BBB; thereby providing new insights that may be relevant for the effective design and delivery of tau-specific immunotherapeutics for the treatment of AD and other tauopathies.

## RESULTS

### APOE4 iBECs demonstrate increased levels of MAPT, tau and p-tau expression compared to APOE3 iBECs

To explore the effects of endogenous tau in BECs in an *APOE* context we utilised *in vitro* BBB models based on *APOE3-* and *APOE4*-carrying human induced pluripotent stem cell (hiPSC)-derived iBECs (*N* = 3 lines per each *APOE* genotype, including one isogenic pair for which *APOE E4/E4* was converted to *iAPOE E3/E3* using CRISPR-Cas9^40^, **Table S1**), a system previously established in our laboratory^41^. We differentiated *APOE3* and *APOE4* hiPSCs into iBECs as validated by the expression of the BBB markers claudin-5 and zonula occludens-1 (ZO-1), and a characteristic cobblestone-like morphology (**Figure 1A**). *APOE3* and *APOE4* expression was validated at the mRNA level using quantitative PCR (qPCR) (**Figure S1A**).

**Figure 1.**
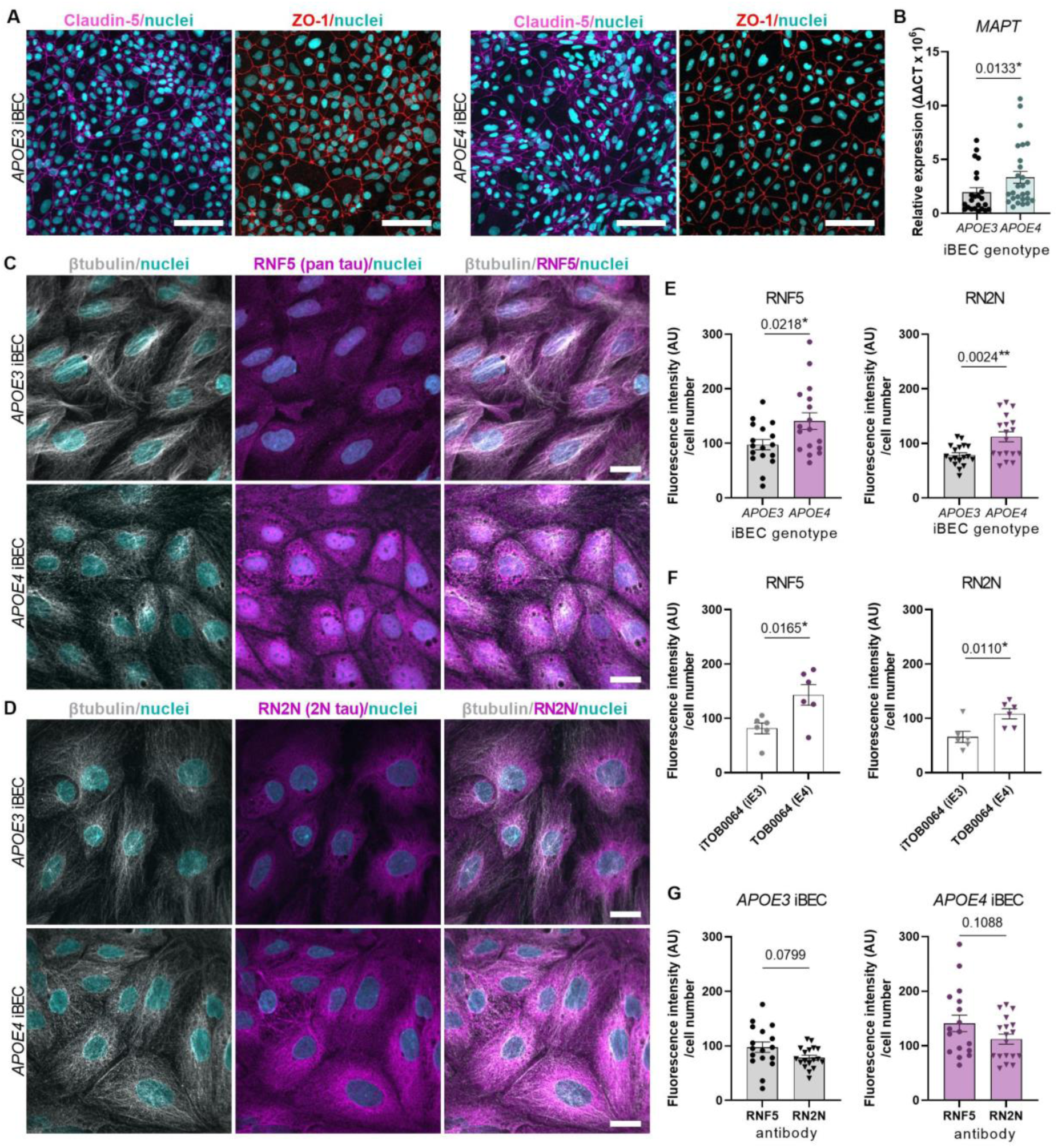
Higher tau expression in *APOE4 compared to APOE3* iBECs. (**A**) Representative immunofluorescence images of claudin-5 (magenta) and ZO-1 (red) with Hoechst nuclear counterstaining (cyan) in *APOE3* and *APOE4* iBECs. Scale bar = 100 μm. (**B**) Relative mRNA expression of *MAPT* in *APOE3* and *APOE4* iBECs. Results presented as ΔΔCT x 10^6^. *N* = 3 biological replicates and a minimum of *n* = 6 independent replicates per line. (**C-D**) Representative high-magnification immunofluorescence images of β-tubulin (grey) and tau detected by anti-pan tau RNF5 antibody (magenta) (C) and anti-2N tau RN2N antibody (magenta) (D) in *APOE3* and *APOE4* iBEC. Nuclei (cyan) stained with Hoechst. Scale bar = 20 µm. (**E**) Fluorescence signal intensity (AU) of RNF5 and RN2N in *APOE3* and *E4* iBEC, normalised to total cell number. *N* = 3 biological replicates and a minimum of *n* = 5 independent replicates per line. (**F**) Fluorescence signal intensity (AU) of RNF5 and RN2N in iTOB0064 (*iAPOE3*) and TOB0064 (*APOE4*) iBECs, normalised to the total cell number. *N* = 1 biological replicate (isogenic pair) and minimum of *n* = 5 independent replicates per line. (**G**) Comparison of fluorescence signal intensity of tau detected with RNF5 and RN2N antibodies in *APOE3* (I) and *APOE4* (J) iBECs, normalised to total cell number. Data analysed with Mann-Whitney *U* test in (B), Student’s *t*-test in (E,F and G:*APOE4* iBEC) and Welch’s *t*-test in (G:*APOE3* iBECs). Error bars = SEM. \**p*<0.05, \*\**p*<0.01.

We then investigated tau expression in iBECs carrying *APOE3* and *APOE4* isoforms and found increased (*p*<0.05) transcription of the microtubule-associated protein tau (*MAPT)* gene, which encodes tau, in *APOE4* cells compared to *APOE3* (**Figure 1B**). Of note, *MAPT* transcription did not differ between the parental *APOE4* line (TOB0064) and its isogenic-corrected *iAPOE3* control line (iTOB0064) (**Figure S1B**). Next, to determine tau protein expression in iBECs we utilised two anti-tau monoclonal antibodies, RNF5 and RN2N, previously developed and extensively characterised by us^42–45^. RNF5 is an IgG2b pan-tau monoclonal antibody specific to all six human tau isoforms^42^, whereas RN2N IgG2a is specific for the human 2N isoform of tau^44^. These antibodies detected expression of tau in both *APOE3* and *APOE4* iBECs by immunofluorescence analysis (**Figure 1C,D**; **Figure S2,S3**), with *APOE4* iBEC presenting higher tau levels than *APOE3* cells recognised by both RNF5 (*p*<0.05) and RN2N (*p*<0.01, **Figure 1E**). To determine whether the observed effect was driven by the *APOE4* allele, we directly compared tau protein expression between the parental TOB0064 (*APOE4*) and isogenic-corrected iTOB0064 (*iAPOE3*) lines and found an increased signal intensity of RNF5 (*p*<0.05) and RN2N (*p*<0.05) in TOB0064 (*APOE4*) compared to iTOB0064 (*iAPOE3*) iBECs (**Figure 1F**). When comparing within iBECs of the same *APOE* genotype, no significant difference was found in the signal intensity corresponding to tau recognised by RNF5 antibody when compared to RN2N (**Figure 1G**) and no signal was found in secondary antibody-only controls for RNF5 and RN2N (**Figure S4**).

Given the role of *APOE4* in aberrant tau phosphorylation^46^, we next assessed tau phosphorylation of epitope Ser396 (p-tau), known to be strongly implicated in AD-related tau pathology^47^. We found p-tau in selected cells in our model (**Figure 2A**) and confirmed their brain endothelial phenotype using co-expression of the BEC marker ZO-1 (**Figure 2B**). Quantification of p-tau revealed an increased (*p*<0.0001) number of iBECs expressing p-tau in monolayers formed by *APOE4*-carrying cells compared to *APOE3* iBECs (**Figure 2C**, **Figure S5**). Similarly, we found an increased (*p*<0.001) number of p-tau positive cells in TOB0064 (*APOE4*) iBEC monolayers compared to isogenic corrected iTOB0064 (*iAPOE3*) cells (**Figure 2D**), confirming that the observed effect is associated with the *APOE4* allele.

**Figure 2.**
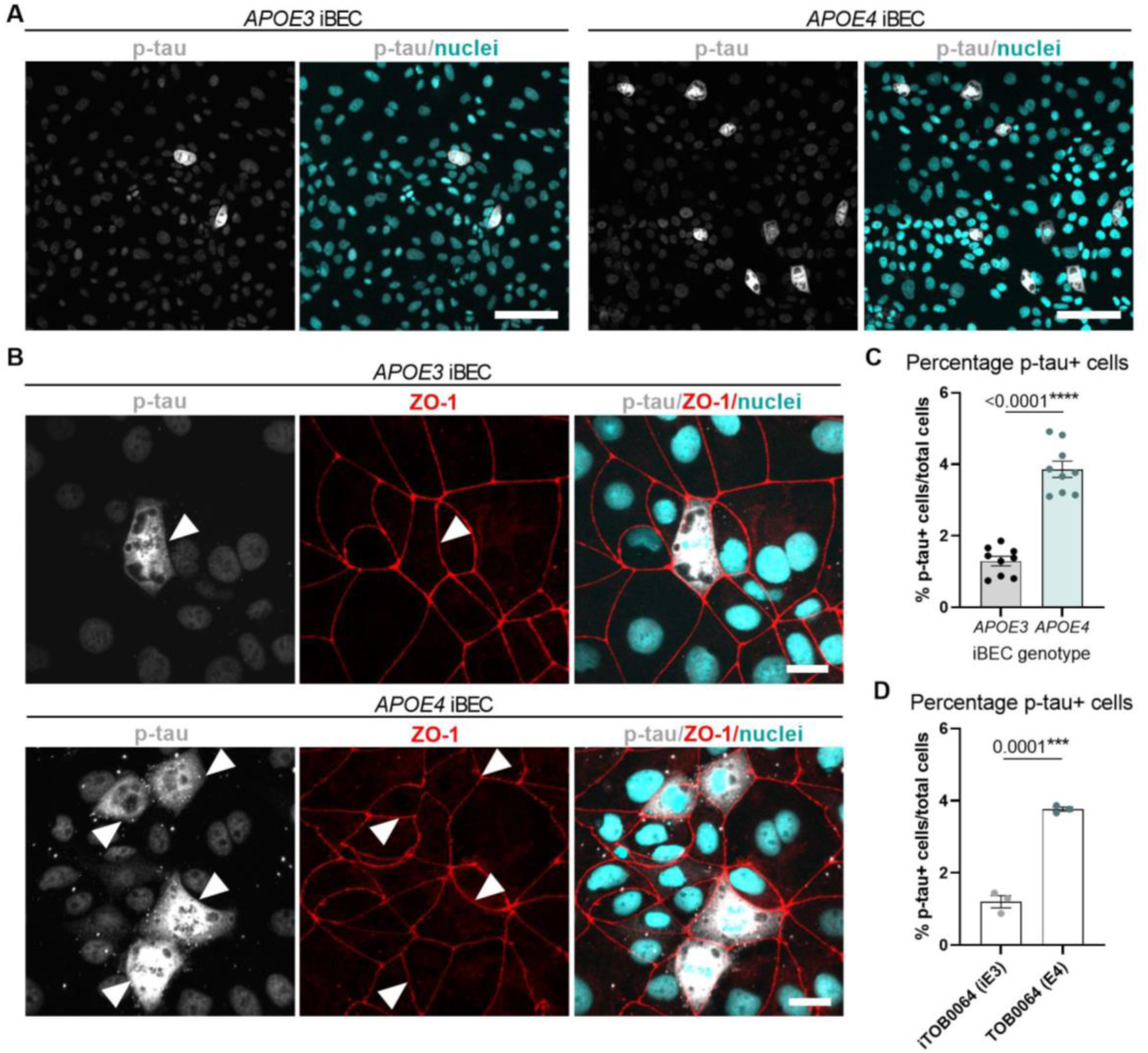
Increased Ser396-phosphorylated tau (p-tau) in *APOE4* compared to *APOE3* iBECs. (**A**) Representative immunofluorescence images of p-tau (grey) with Hoechst nuclear counterstaining (cyan) in *APOE3* and *APOE4* iBECs. Scale bar = 100 μm. (**B**) Representative high-magnification immunofluorescence images of p-tau positive cells (grey) and ZO-1 (red) in *APOE3* and *E4* iBECs. Nuclei (cyan) stained with Hoechst. Scale bar = 20 µm. (**C**) Quantification of p-tau-positive cells in monolayers formed by *APOE3* and *APOE4* iBECs. Data presented as percent of p-tau-positive cells normalised to total cell number. *N* = 3 biological replicates and minimum of *n* = 3 independent replicates per line. (**D**) Quantification of p-tau-positive cells in monolayers formed by iTOB0064 (*iAPOE3*) and TOB0064 (*APOE4*) iBEC. *N* = 1 biological replicate (isogenic pair) with *n* = 3 independent replicates per line. Data analysed with Student’s *t*-test in (C,D). Error bars = SEM. \*\*\**p*<0.001, \*\*\*\**p*<0.0001.

### Passive permeability of anti-tau therapeutic antibodies is reduced in APOE4 iBECs

Insufficient transport at the BBB is a known hurdle for the development of large-molecule immunotherapeutics^48^. Hence we hypothesised that increased levels of tau in *APOE4* iBECs may contribute to limited anti-antibody passage through the BBB. To investigate this, we further utilised the tau-targeting antibodies RNF5 and RN2N shown to achieve therapeutic effects in tau transgenic mouse models^42–44^ and screened their permeability *in vitro* in our Transwell-based *APOE* BBB model^41^.

We previously demonstrated that Transwell inserts with 0.4 μm pores, most commonly used in the iPSC-BBB models^34,49–52^, are suitable to assess small molecule (5 kDa) permeability, while those with pores of 3.0 μm in diameter provide a more adequate *in vitro* system to study larger (150 kDa) molecule transport^41^. As demonstrated by us previously^41^ and in this study, when cultured in Transwell inserts containing membrane pores with a diameter of either 0.4 μm or 3.0 μm, *APOE4* iBEC formed monolayers of reduced TEER (*p*<0.0001) compared to *APOE3* controls (0.4 μm Transwells: *APOE3* 3018 ± 75.19 Ohm x cm^2^, *APOE4* 1608 ± 95.31 Ohm x cm^2^; 3.0 μm Transwells: *APOE3* 1716 ± 39.37 Ohm x cm^2^, *APOE4* 817.6 ± 85.42 Ohm x cm^2^, mean ± SEM, **Figure 3A,B**). This corresponded to the TEER values of the BBB previously found *in vivo* (average of 1462 Ohm x cm^2^ in^53^ and 1870 Ohm x cm^2^ in^54^). We also found increased permeability to biologically inert fluorescent FITC-conjugated 5 kDa and 150 kDa tracers in *APOE4* iBEC monolayers compared to *APOE3* iBEC when cultured in respective 0.4 μm and 3.0 μm pore Transwells (*p*<0.0001, **Figure 3C,D**), suggesting the presence of a clinically relevant^28^ ‘leaky BBB’ phenotype in *APOE4-*carrying cells. We then evaluated the passive permeability of the anti-tau antibodies RNF5 and RN2N (∼150 kDa in size) in the *APOE3* and *APOE4* iBEC Transwell model (3.0 μm pore) where the tested antibody was added to the luminal (top) chamber of a Transwell at 1 μM and its concentration in the abluminal (bottom) chamber was assessed at 24 h as we previously established^41^, utilising an ELISA-based detection of antibodies in the Transwell flow-through. This revealed a striking decrease in passive transport of both RNF5 (*p*<0.01) and RN2N (*p*<0.001) through monolayers formed by *APOE4* iBECs compared to *APOE3* iBECs (**Figure 3E**). Similarly, a decrease in RNF5 (*p*<0.0001) and RN2N (*p*<0.0001) permeability was observed in TOB0064 (*APOE4*) when compared to its isogenic corrected control iTOB0064 (*iAPOE3*) iBEC (**Figure 3F**), despite TOB0064 cells forming a barrier of reduced (*p*<0.0001) integrity as compared to iTOB0064 (**Figure S6**). Together, this suggests a strong link between reduced anti-tau antibody permeability and the *APOE4* allele. When comparing within cells of the same *APOE* genotype, we also observed increased passive permeability of RN2N as opposed to RNF5 (RNF5 vs RN2N in *APOE3* iBECs *p*<0.05, in *APOE4* iBECs *p*<0.01, **Figure 3G**), possibly reflecting the selective binding of RN2N to 2N tau, that comprises only a fraction of total tau^44^. This reduced passive leakage was also seen in *APOE4* iBECs for AlexaFluor647 (AF647)-conjugated RNF5 and RN2N by measuring antibody-associated AF647 fluorescence in media collected from the bottom chamber of the Transwell system (*p*<0.05, **Figure S7A**). Intriguingly, we did not observe any differences in the passive permeability of the AF647-conjugated anti-amyloid antibody Aducanumab between *APOE3* and *APOE4* cells, suggesting the observed effect may be specific to tau-targeting antibodies (**Figure S7A**). Observation of similar effects of *APOE4* on RNF5 and RN2N permeability, when assessed with both ELISA (**Figure 3E**) and fluorescence-based methods (**Figure S7A**), also confirmed that the addition of a fluorescent tag to the antibody did not affect its permeability dynamics in iBECs. To further confirm anti-tau antibody permeability and *APOE4* association, we employed our alternative, previously characterised iBEC model comprising familial AD patient-derived iBECs carrying the disease-associated mutation (exon 9 deletion, *ΔE9*) in the presenilin-1 (*PSEN1*) gene with those cells being simultaneously *APOE3* carriers^50,55^. Interestingly, the passive permeability of AF647-RN2N and AF647-Aducanumab did not vary between controls (*PSEN1* wildtype, *APOE3*) and familial AD (*PSEN1-ΔE9*, *APOE3*) iBECs suggesting the dependence of tau antibody permeability on the presence of *APOE4* allele, rather than an AD-related phenotype more generally (**Figure S7B**).

**Figure 3.**
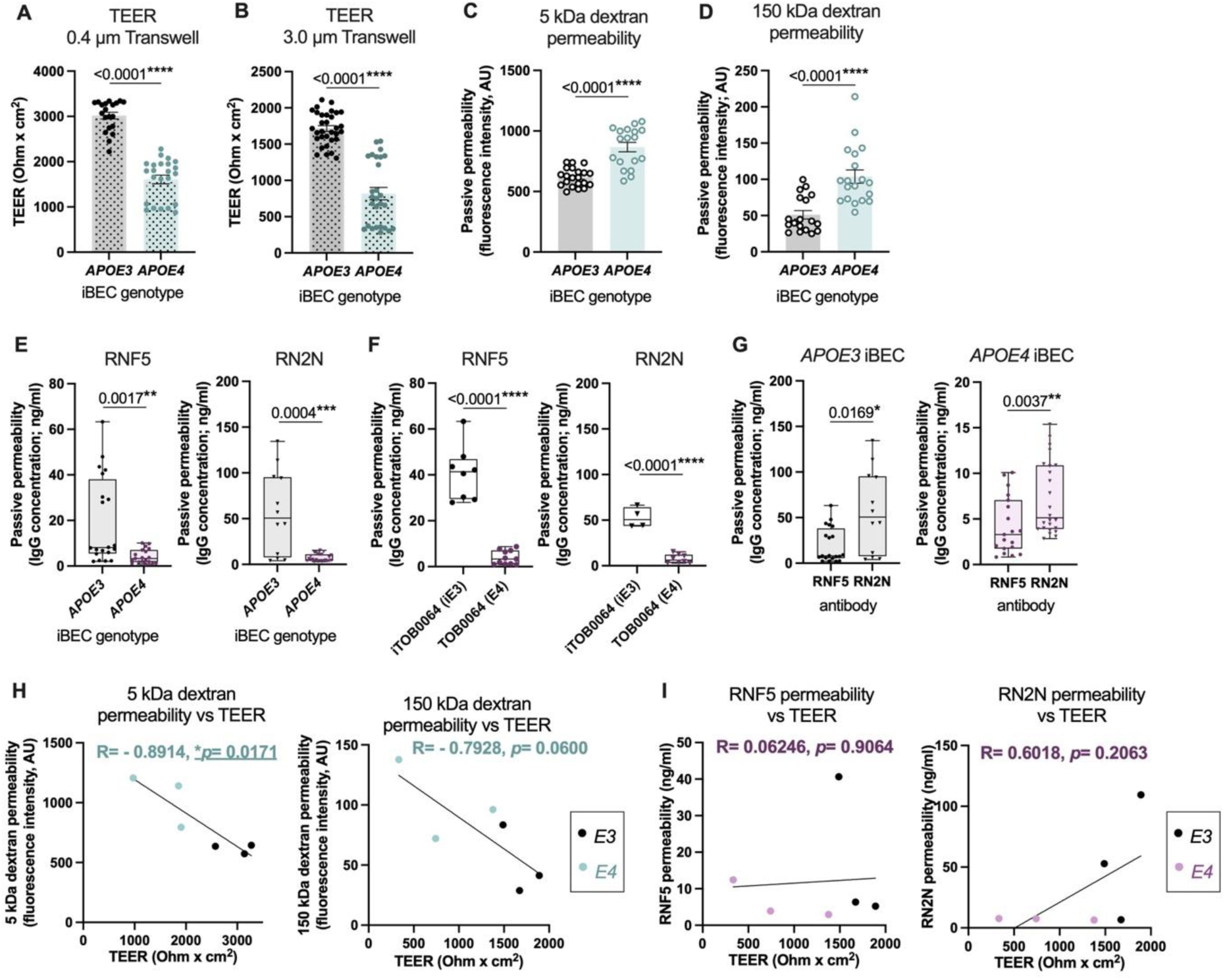
Barrier properties and reduced passive permeability of anti-tau antibodies RNF5 and RN2N in *APOE4* iBECs. (**A-B**) Trans-endothelial electrical resistance (TEER, shown as Ohm x cm^2^) of *APOE3* and *APOE4* iBEC, measured in Ø 0.4 µm and Ø 3.0 µm pore Transwells. *N* = 3 biological replicates and minimum *n* = 6 independent replicates per line. (**C-D**) Passive permeability of 5 kDa and 150 kDa FITC-conjugated dextran in *APOE3* and *APOE4* iBECs. Data presented as a fluorescence intensity (AU) of FITC-conjugated dextran measured in the bottom chamber of the Transwell at 24 h. *N* = 3 biological replicates and minimum *n* = 5 independent replicates per line. (**E**) Passive permeability RNF5 and RN2N antibodies in *APOE3* and *APOE4* iBECs. Data presented as an antibody concentration (ng/ml) detected in the bottom chamber of the Transwell at 24 h. *N* = 3 biological replicates and minimum *n* = 2 independent replicates per line. (**F**) Passive permeability of RNF5 and RN2N in iTOB0064 (*iAPOE3*) and TOB0064 (*APOE4*) iBEC. *N* = 1 biological replicate (isogenic pair) and a minimum of *n* = 4 independent replicates per line. (**G**) Comparison between RNF5 and RN2N passive permeability in *APOE3* and *APOE4* iBECs. *N* = 3 biological replicates and minimum *n* = 2 independent replicates per line. (**H**) Correlation between averaged (per line) *APOE* iBEC barrier integrity (TEER) and averaged (per line) passive permeability of 5 kDa dextran or 150 kDa dextran. *N* = 3 biological replicates. (**I**) Correlation between averaged (per line) *APOE* iBEC barrier integrity (TEER, Ø 3.0 µm pore Transwells) and averaged (per line) passive permeability RNF5 and RN2N antibodies. *N* = 3 biological replicates. Data analysed with Mann-Whitney *U* test in (A-E, G) and Student’s *t*-test in (F) and Pearson’s correlation in (H-I). Error bars = SEM in bar graphs. Whiskers = min-max in box plots. \**p*<0.05, \*\**p*<0.01, \*\*\**p*<0.001, \*\*\*\**p*<0.0001.

Since the barrier in our *APOE4* model was seemingly ‘more leaky’ (**Figure 3A-D**) while simultaneously not facilitating increased permeability of anti-tau antibodies (**Figure 3E**), we further investigated the correlation between cargo permeability and barrier integrity in *APOE* iBECs. When considering the biologically inert FITC-conjugated dextran tracers, as expected, we observed a negative correlation between iBEC barrier integrity (TEER) and its permeability, with lower barrier integrity correlating with higher permeability to both small molecule (5 kDa) dextran (Pearson’s R= −0.8914, *p*<0.05) and larger molecule (150 kDa) dextran (Pearson’s R= −0.7928, *p*= 0.06) as assessed in 0.4 μm and 3.0 μm Transwell formats, respectively (**Figure 3H**). This dependence, however, was not replicated for anti-tau antibodies where RNF5 passive permeability largely did not correlate with iBEC TEER (Pearson’s R= 0.06246, *p*= 0.9064) while RN2N showed an opposite trend to that observed for dextrans permeability (Pearson’s R= 0.6018, *p*= 0.2063, (**Figure 3I**)).

Given that previously detected tau expression in *APOE* iBECs (**Figure 1C,D**), we next evaluated iBEC monolayers cultured on Transwells in the presence of AF647-conjugated RNF5 and RN2N for 24 h and detected a fluorescence signal in *APOE3* and *APOE4* iBECs, suggesting that the antibody is, at least in part, being trapped in the cytoplasm during passage between the top and bottom chambers of the Transwell system (**Figure 4A,B**, **Figure S8**). To confirm specificity for tau antibodies and not just large molecules in general, we exposed cells grown on Transwell inserts to 150 kDa FITC-conjugated dextran for 24 h, however, did not detect any robust signal of trapping (**Figure 4C**, **Figure S8**). Finally, to confirm that the observed effects were due to anti-tau antibodies potentially recognising their target in *APOE4* iBECs rather than IgG generally binding to cells, we compared the passive permeability of RNF5, RN2N and their respective IgG2b and IgG2a controls in *APOE* iBECs. As expected given the moderate levels of total and 2N tau in *APOE3* iBECs as compared to *APOE4* iBEC (**Figure 1E**), the passive permeability of RNF5 and RN2N did not significantly differ from their respective IgG controls in *APOE3* cells (**Figure 4D,E**). However, in *APOE4* iBECs, control IgGs showed significantly increased passive permeability compared to RNF5 (*p*<0.0001) and RN2N (*p*<0.0001, **Figure 4D,E**) pointing to tau-specific interactions of RNF5 and RN2N antibodies in *APOE4* iBECs.

**Figure 4.**
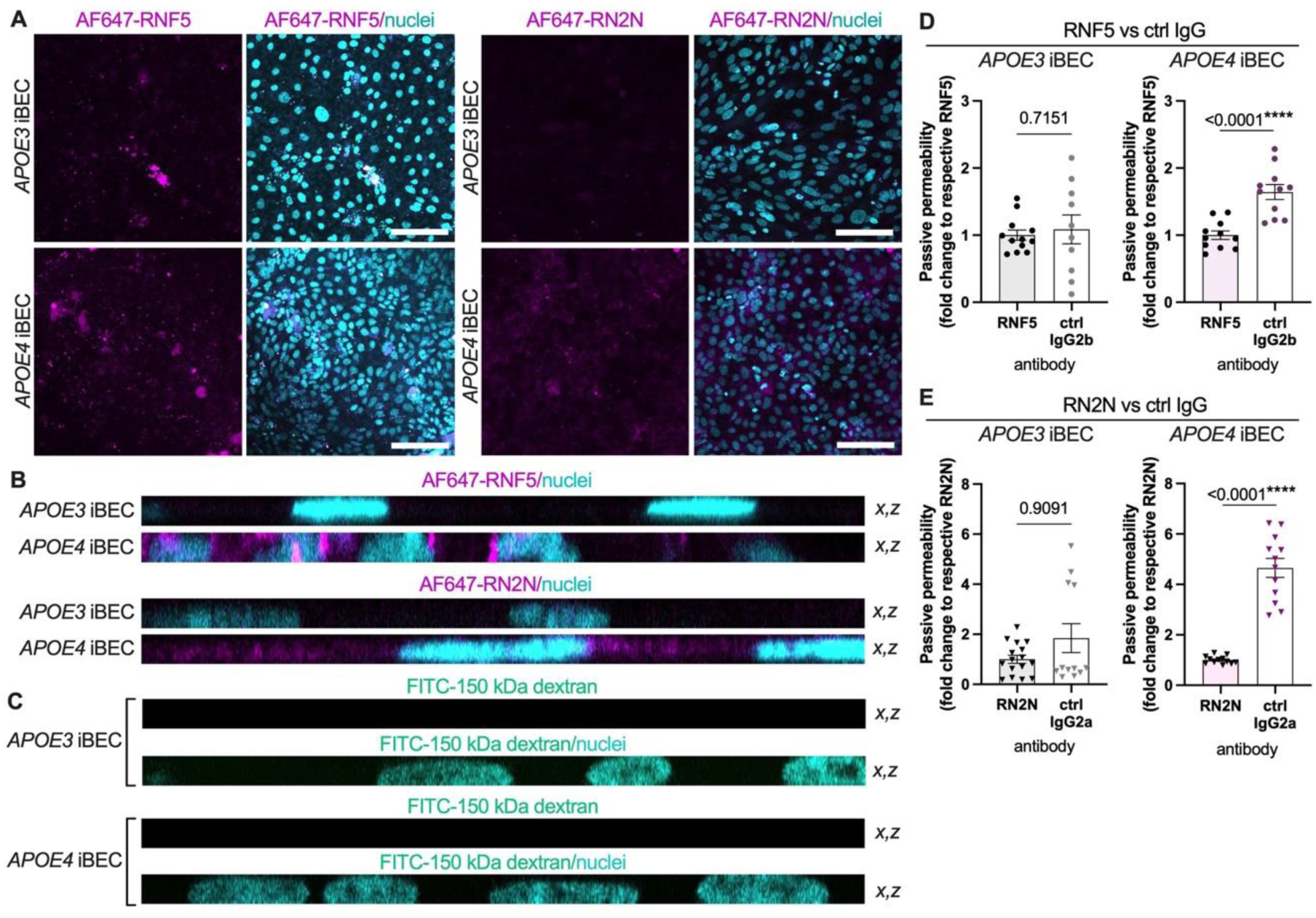
Presence of RNF5 and RN2N in *APOE* iBEC monolayers and their reduced permeability compared to isotype controls in *APOE4* cells. (A) Detection of AF647-RNF5 and AF647-RN2N signal (magenta) in *APOE3* and *APOE4* iBEC monolayers cultured on Transwells, at 24 h post-treatment with the respective antibody. Nuclei counterstained with Hoechst (cyan). Scale bar = 100 μm. (**B-C**) Orthogonal (x,z) views of *APOE* iBEC monolayers exposed to AF647-RNF5 (magenta), AF647-RN2N (magenta) and FITC-conjugated 150 kDa dextran (green) for 24 h. Nuclei counterstained with Hoechst (cyan). (**D-E**) Comparison of passive permeability of RNF5 and RN2N to respective IgG controls in *APOE3* and *APOE4* iBECs. Data presented as fold changes in control IgG concentration (ng/ml) detected in the bottom chamber of the Transwell at 24 h, normalised to respective anti-tau antibodies within each line. *N* = 3 biological replicates and minimum *n* = 2 independent replicates per line. Data in (D) were analysed with Welch’s *t-*test (*APOE3*) and Student’s *t-*test (*APOE4*). Data in (E) were analysed with Mann-Whitney *U* test (*APOE3*) and Welch’s *t*-test (*APOE4*). Error bars = SEM. \*\*\*\**p*<0.0001.

Together, these observations suggest that elevated levels of tau in *APOE4*-carrying iBECs may contribute to the reduced permeability of tau-targeting RNF5 and RN2N antibodies in the *in vitro* BBB system.

### Active delivery with FUS^+MB^ enhances the permeability of tau immunotherapeutics across APOE4 iBECs

Having observed reduced passive permeability of anti-tau antibodies in *APOE4* cells, we hypothesised that active delivery methods may be required to facilitate the delivery of tau-targeting immunotherapeutics in *APOE4* iBECs. Focused ultrasound and microbubble (FUS^+MB^) technology is an emerging method successfully utilised to enhance large molecule drug delivery in preclinical AD studies performed by us and others^42,44,45,56–58^ and proven safe and clinically effective in temporary BBB opening in AD patients^57,59–62^. We have previously established an ultrasound-mediated antibody delivery *in vitro* platform where we demonstrated that FITC-conjugated 150 kDa dextran, AF647-RNF5 and AF647-Aducanumab permeability can be increased in a sporadic AD BBB model by the application of FUS^+MB^ at optimised parameters^41^. Here we reproduced FUS^+MB^-enhanced RNF5 antibody delivery in our model^41^ and further trialled the utility of FUS^+MB^ to deliver RN2N and control IgGs in *APOE3* and *APOE4* iBEC. Using ELISA-based antibody detection, we found FUS^+MB^ treatment improved the passage of RNF5 antibody in both *APOE3* (*p*<0.01) and *APOE4* (*p*<0.001) iBECs (**Figure 5A**). Similarly, FUS^+MB^ improved delivery of RNF5’s control IgG2b (*p*<0.01). However, the delivery efficiency was higher (*p*<0.05) for control IgG compared to RNF5 in *APOE4* cells while showing similar delivery efficiency in *APOE3* cells (**Figure 5A**) suggesting interactions of RNF5 with iBECs could affect its delivery with FUS^+MB^ in *APOE4* cells. FUS^+MB^ also effectively improved the permeability of RN2N (*APOE3* iBECs: *p*<0.0001, *APOE4* iBECs: *p*<0.01) and its corresponding control IgG2a (*APOE3* iBEC: *p*<0.05, *APOE4* iBEC: *p*<0.01), while reaching similar delivery efficiency when comparing both tau-specific and control IgG antibodies formats (**Figure 5B**). Finally, we detected no difference in the delivery efficiency of RN2N compared to RNF5 in *APOE* iBECs following FUS^+MB^ (**Figure 5C**).

**Figure 5.**
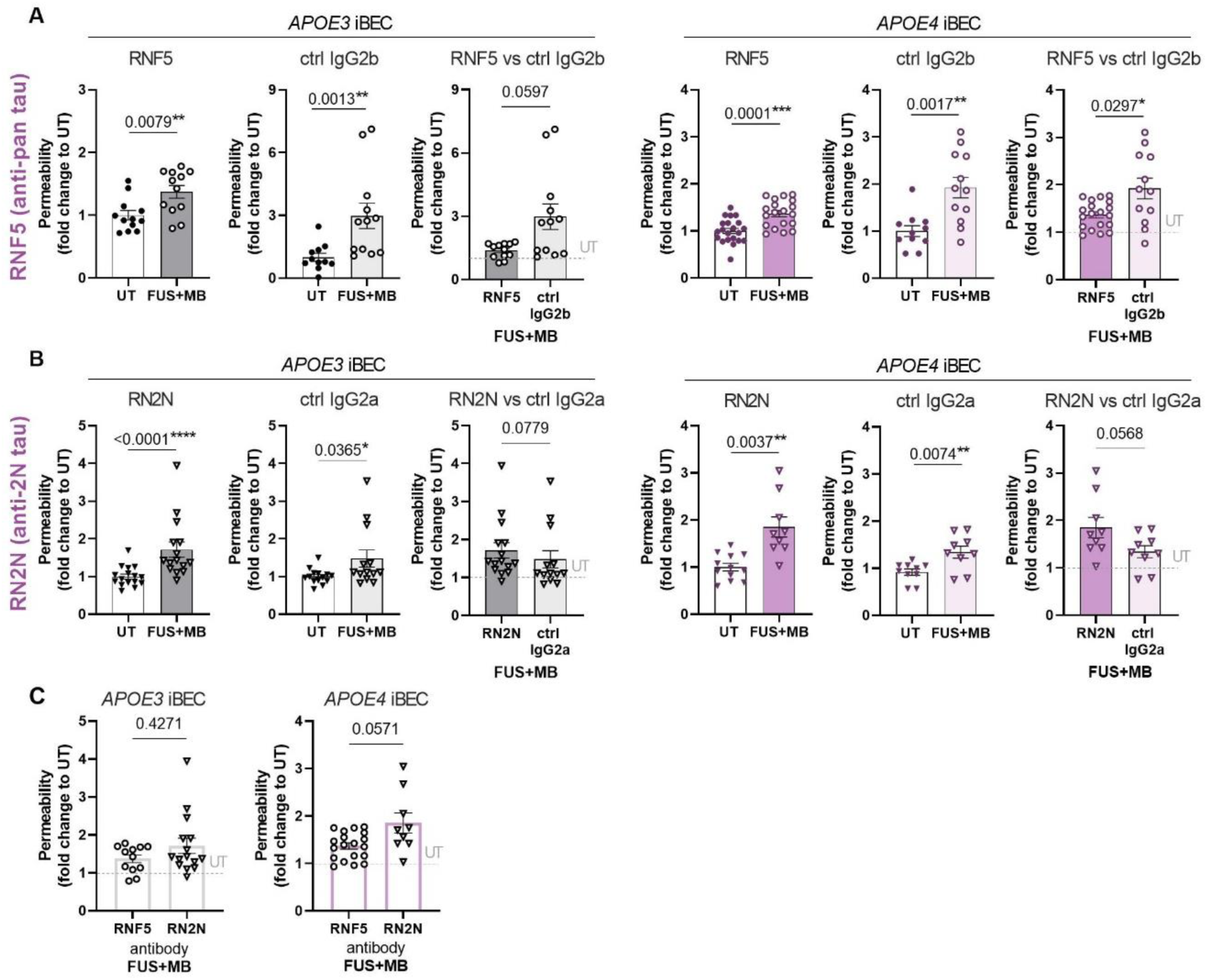
Focused ultrasound and microbubble (FUS^+MB^) technology aid in tau-immunotherapeutics delivery in *APOE* iBEC. (**A**) FUS^+MB^ mediated delivery of RNF5 and its corresponding control IgG in *APOE3* and *APOE4* iBEC. (**B**) FUS^+MB^ mediated delivery of RN2N and its corresponding control IgG in *APOE3* and *APOE4* iBEC. (**C**) Comparison of FUS^+MB^-mediated delivery efficiency of RNF5 and RN2N between *APOE3* and *APOE4* iBEC. Data presented as fold changes in anti-tau antibody or control IgG concentration (ng/ml) detected in the bottom chamber of the FUS^+MB^-exposed Transwell at 24 h, normalised to respective untreated (UT) control within each line. In (A-C): *N* = 3 biological replicates and minimum *n* = 2 independent replicates per line. Data analysed with Student’s *t*-test, Welch’s *t*-test or Mann-Whitney *U* test as appropriate, error bars = SEM. \**p*<0.05. \*\**p*<0.01, \*\*\**p*<0.001.

## DISCUSSION

Tau-specific immunotherapy is emerging as a promising treatment approach for AD in preclinical studies^44,63–68^ with various early-stage clinical trials currently ongoing^2,69^. However, previous studies have revealed that achieving a sufficient antibody concentration in the brain is challenging^70,71^. Therefore, to improve the delivery of tau immunotherapeutics into the patient’s CNS, it is crucial to understand how the BBB cells interact with the drug and elucidate molecular and cellular mechanisms that limit therapeutic antibody passage through the BBB.

Here our goal was to investigate the interactions of anti-tau therapeutic antibodies with human BECs *in vitro* and identify cell-type-specific effects that could influence BBB permeability of tau-targeting immunotherapeutics in AD. By utilising our hiPSC-derived sporadic AD BBB model^41^ and two tau-specific monoclonal antibodies, RNF5 and RN2N^42–45^, we found increased endogenous tau expression and Ser396 phosphorylation to be associated with the *APOE4* allele in iBEC, suggesting that as for other brain cell-types^22–25,72,73^, *APOE4* may exacerbate tau-related changes in BECs. This tau expression in *APOE4* iBECs was further associated with decreased passive permeability of the anti-tau therapeutic antibodies RNF5 and RN2N across the BBB *in vitro*. We also identified an accumulation of tau antibodies within the *APOE4* iBEC monolayer, together pointing to cell-type-specific molecular interactions within BECs that could limit tau immunotherapeutics delivery across the BBB, hence minimizing their target engagement within the brain parenchyma. We also found that barrier integrity as measured by TEER is a read-out for the permeability rate of biologically inert fluorescent dextran tracers, while the same observation was not made for anti-tau antibodies. This may suggest that for certain therapeutics the molecular interactions with the BBB cells need to be considered in addition to actual barrier integrity, potentially explaining the lack of widespread drug leakage through the BBB despite the barrier’s breakdown in AD^37,38^. With further investigation required, our data may also provide indirect evidence for dextran tracers to permeate through the iBEC monolayer by the paracellular route while antibodies, at least in part, enter via the transcellular pathway, as has been previously proposed for the IgG transport at brain barriers^74^. Moreover, we found the limited permeability of RN2N and RNF5 in *APOE4* iBECs could be partially overcome by applying FUS^+MB^ ^39^, suggesting passive immunotherapy on its own may not be the most effective avenue to tau-targeted drug delivery and instead, the development of active-delivery technologies may be needed to realise the clinical potential of tau immunotherapies in *APOE4* individuals.

With more studies required to fully decipher the molecular interactions between tau and *APOE4* at the BBB, here we postulate that the *APOE4* allele may influence the expression and phosphorylation of tau in BECs, which may contribute to pathological changes at the BBB and affect anti-tau therapeutic antibody delivery in sporadic AD.

A few independent lines of evidence suggests BEC involvement in APOE4-or tau-related BBB dysfunction^14–16,26,75–78^. However, with both proteins being expressed in multiple brain cell types, it remains challenging to dissect their cell-type-specific contributions to BBB pathophysiology in AD. By utilising human *APOE3* and *APOE4* iBEC models to investigate tau expression in a single cell type system, we identified an increase in tau protein levels and its phosphorylation in *APOE4* iBECs to be associated with *in vitro* BBB dysfunction. Although direct causality is yet to be determined, this may suggest the potential involvement of endogenous tau within BECs in the cerebrovascular dyshomeostasis observed in *APOE4* carriers^28^. Interestingly, *APOE4* hiPSCs utilised in our study were generated from presymptomatic *APOE4* donors^40^, indicating that tau-related changes in BECs could occur early and precede the cognitive decline in *APOE4*-bearing individuals, corroborating previously reported early BBB dysfunction in cognitively normal *APOE4* carriers^28^. This further suggests our model may be suitable for studying BBB impairment prevention strategies for sporadic AD.

Although the molecular mechanism of *APOE4* and tau interaction was not investigated in this model, an interactome map of S396 and S404 p-tau in the human AD brain identified APOE as one of the proteins directly mediating pathological actions of p-tau in neuronal neurofibrillary tangles, suggesting an intimate molecular crosstalk between APOE and tau in AD^47^. Interestingly, in our study, we identified increased transcription of tau-encoding *MAPT* gene in *APOE4* iBECs, while this effect was not found when comparing parental TOB0064 (*APOE4*) and isogenic corrected iTOB0064 (*iAPOE3*) lines. Simultaneously, tau protein expression and phosphorylation were consistently increased in TOB0064 (*APOE4*) cells compared to iTOB0064 (*iAPOE3*), suggesting that *APOE4* may have a more discernible effect on tau protein expression than *MAPT* gene transcription in iBECs. Alternatively, increased levels of intracellular tau in *APOE4* iBECs may in part result from impaired tau clearance or ineffective protein degradation pathways in these cells, with the exact mechanism remaining to be elucidated.

Increased levels of endogenous tau in *APOE4* iBECs identified herein were also associated with limited passive permeability of tau-specific therapeutic antibodies at the *in vitro* barrier. Intriguingly, this effect was solely observed for tau-targeting antibodies as we did not detect any differences in the passive permeability of the anti-amyloid antibody Aducanumab (Aduhelm^56,79^) between *APOE3* and *APOE4* iBECs. Notably, our model did not incorporate vessel-associated amyloid^80,81^, and therefore may not fully represent an AD-like BBB environment. However, another study utilising *APOE3* and *APOE4* iBECs previously detected *APOE4*-dependent increased production of amyloid-β 40 and 42 species by iBECs^82^, hence it cannot be excluded that endogenous amyloid-β is present and physiologically influences antibody permeability in our model system. The limited permeability of tau antibodies through the *in vitro* BBB was also strongly associated with the *APOE4* genotype as experiments conducted with our familial AD patient-derived model^50,55^ comprising iBECs carrying the *PSEN1-ΔE9* mutation and simultaneously the *APOE3* alleles, did not reveal any significant differences in passive permeability of anti-tau antibody or Aducanumab compared to respective control cells. While in addition to tau expression, other mechanisms such as altered vesicular transport^8,83,84^ or transporters expression^84^ may contribute to observed effects in *APOE4* cells, our observations highlight that careful assessment of AD patient’s genetic profile may be critical for the successful delivery of immunotherapies at the level of their brain barriers, especially in *APOE4* carriers who comprise over half of the AD patient population^85^.

Our results suggest that increased intracellular accumulation of tau in *APOE4* BEC and consequent molecular interaction with tau-specific immunotherapeutics within BECs may represent one of the mechanisms by which the passive delivery of anti-tau antibodies at BBB is compromised. Thus active tau immunotherapeutic delivery methods such as FUS^+MB^ ^57,59–62^ may be required to enhance the transport of tau immunotherapeutics at otherwise ‘leaky’ BBB of *APOE4*-carrying patients. Here, similar to our previous *in vivo*^42,44,45^ and *in vitro*^41^ studies, we successfully enhanced therapeutic tau antibody transport in the sporadic AD BBB model, further validating the utility of FUS^+MB^ technology in antibody delivery in AD. We found lower delivery efficiency of RNF5 compared to its non-tau binding IgG isotype control in *APOE4* iBECs, suggesting certain anti-tau antibodies may need additional modifications such as encapsulation in nanoparticle, liposomes or exosomes^48^ to effectively permeate through BBB in *APOE4* carriers. This asks for the careful design of novel tau-targeting immunotherapeutics factoring in the antibody’s therapeutic effects in the brain parenchyma as well as its molecular interactions with BEC that could significantly limit its BBB permeability.

With our model recapitulating aspects of the clinical BBB phenotype observed in both *APOE4* carriers^28^ as well as tauopathies^12,15,16,86^, and drug permeability dynamics assessed in hiPSC-derived iBEC platforms correlating with that of human BBB^87,88^, we expect our results to be of high translational relevance, and upon validation in the *in vivo* systems, prove important for tau immunotherapy delivery in *APOE4* human carriers. Yet, further studies are needed to comprehensively understand the association and causality among *APOE* genotype, tau, BBB function and the permeability of tau-targeting therapeutics in AD, potentially via novel *in vitro* and *in vivo* models. Currently, tau overexpression is driven by neuron-specific promoters in the majority of transgenic mouse models^16,18,21,42,44,45^. Ideally, developing novel mouse models expressing tau under BEC-specific promoters, such as Tie2^89^, would shed light on the contribution of tau to neurovascular unit pathology and further expand understanding of the role of non-neuronal tau in AD. Similarly, novel hiPSC lines carrying disease-associated *MAPT* mutations^90–92^, *APOE4* knockout/knockin sequences, or those expressing tau aggregation biosensors^93^ could be utilised to develop more complex, multicellular^34^ hiPSC-derived *in vitro* BBB models and aid in in-depth characterisation of tau effects on the BEC phenotype and function. Additionally, given the known inter-cell line variability of hiPSCs^40,41,94^, future studies incorporating larger patient cell cohorts and validation in human brain samples would be needed to confirm the results presented here. The former paired with drug screening of multiple formats of anti-tau therapeutic antibodies could effectively inform drug formulation design in tau immunotherapy and lead to the identification of the most promising drug candidates. Furthermore, exploring avenues to reduce tau levels in BECs could aid in regulating some of the early BBB pathology seen in *APOE4* carriers. Since constitutive tau expression is required for adequate cell function, *APOE4* rather than tau could be a potential therapeutic target in BECs, with its expression altered via immunotherapy^95^, cell type-specific AAV-CRISPR-mediated gene therapy^96^ or ligand-conjugated anti-sense oligonucleotides (ASOs)^97,98^. With BECs being the only brain cells directly and easily accessible to peripherally administered therapeutics, developing such strategies may prove useful in diminishing the detrimental effects of *APOE4* and tau on the BBB, and ultimately, cognition^8,9,28^.

Together, by uncovering the link between the *APOE4* and increased intracellular accumulation and abnormal phosphorylation of tau, and consequently decreased anti-tau therapeutics delivery in patient-derived iBEC, our study expands the understanding of the role of tau at the BBB in AD and paves the way for more effective design of therapeutics targeting tau-associated neurodegeneration.

## MATERIALS AND METHODS

### Human iPSC culture and differentiation towards induced brain endothelial-like cells (iBECs)

Previously published and characterised *APOE3*- and *APOE4*-carrying hiPSC lines (*N* = 3 *APOE3* lines including one isogenic corrected line converted from *APOE4* to *iAPOE3*, and *N* = 3 *APOE4* lines) were used to generate induced brain endothelial-like cells (iBECs), as previously described by us^41^ (**Table S1**). Selected experiments also involved our familial AD model, based on hiPSCs carrying *PSEN1-ΔE9* mutation (*N* = 2 lines) and their corresponding isogenic-corrected and healthy donor controls (*N* = 3 lines), previously developed and characterised by us^50,55^ (**Table S1**). hiPSCs were cultured on human recombinant vitronectin in StemFlex^TM^ media (Life Technologies) and differentiation was initiated by plating singularised hiPSCs on human embryonic stem cells (hESC)-qualified Matrigel (Corning) coating in StemFlex^TM^ media supplemented with 10 μM Rho-associated kinase inhibitor (iROCK)^41,50^. hiPSC were then spontaneously co-differentiated in unconditioned media consisting of DMEM/F12+GlutaMAX (Life Technologies), 20% KnockOUT serum replacement (Life Technologies), 1 x non-essential amino acids (Life Technologies) and 0.1 mM β-mercaptoethanol (Sigma)^41,49,50^. Following six days in unconditioned media, culture media was replaced with endothelial cell media (EC; Life Technologies) supplemented with 2% B27 (Life Technologies), 20 ng/ml basic fibroblast growth factor (FGFb; Peprotech) and 10 μM retinoic acid (RA)^41,49,50^. After two days, cells were purified on collagen IV from human placenta (Sigma) and human plasma fibronectin (Life Technologies) coated plastic culture plates or Ø 0.4 μm or Ø 3.0 μm pore Transwell inserts (Corning) as we described previously^41,55^. The Transwell insert pore diameters of 0.4 μm and 3.0 μm were previously established to be suitable for assessing *in vitro* permeability of small (5 kDa dextran) and large (150 kDa dextran, antibody) molecules, respectively^41^. Following approximately 24 h, cell media was replaced with EC+B27 and cells were allowed to mature for one additional day before performing the assays. iBECs were cultured under normoxia (37°C, 5% CO_2_) conditions in EC+B27 for the duration of performed assays^41,49,50^.

### iBECs phenotype and barrier integrity characterisation

Generated iBECs were characterised for expression of BEC-specific markers ZO-1 and claudin-5 by immunofluorescence as we previously described^41^. Briefly, cells were fixed with 4% paraformaldehyde (PFA) for 15 min at room temperature (RT), permeabilised with 0.3% Triton X-100 for 10 min and blocked for 1 h at RT with 2% bovine serum albumin (BSA, Sigma)/2% normal goat serum (GS, Chemicon) in PBS and incubated with primary antibodies for claudin-5 and ZO-1 (**Table S2**) diluted at 1:100 in a blocking solution overnight at 4°C. The next day, cells were washed with PBS and incubated with secondary antibodies (AlexaFluor-488 or AlexaFluor-647; **Table S2**) diluted at 1:250 in a blocking solution for 1 h at RT in the dark. Cells were washed with PBS and Hoechst (1:5000) counterstain was performed. The coverslips with cells were mounted with Dako Mounting Medium (Agilent). Images were obtained at 20x magnification using a Zeiss LSM-780 confocal microscope.

iBEC barrier integrity was assessed by measuring transendothelial electrical resistance (TEER) across iBEC monolayer using the EVOM3 Volt/Ohmmeter (World Precision Instruments) in 24-well, 6.5 mm Transwell with 0.4 μm pore or 3.0 μm pore membrane insert (Corning) as we previously described^41^. TEER was then measured in three areas per Transwell and the resistance of the blank (no-cells) Transwell was subtracted before averaging. The resulting value was multiplied by the surface area of the Transwell membrane (0.33 cm^2^) for calculation of the final TEER values (Ohm x cm^2^).

### Tau and p-tau immunofluorescence analysis

For immunofluorescence-based quantification of tau expression and tau phosphorylation in iBEC, cells were cultured on collagen IV and fibronectin-coated plastic coverslips under normal conditions, fixed with 4% PFA for 15 min at RT, and immunostaining for the particular marker was performed as described above.

For tau assessment, monoclonal tau-specific antibodies RNF5 (anti-pan tau) and RN2N (anti-2N tau) were generated in-house as previously described^42–44^ and utilised as primary antibodies at optimised 0.05 mg/ml concentration. Additional samples were co-stained with commercially sourced anti-β-tubulin antibody (1:100) (**Table S2**). Anti-Ser396-phosphorylated tau primary antibody was commercially sourced and used at 1:100 (**Table S2**). Antibody specificity was confirmed by performing secondary antibody-only controls.

Following immunostaining, the coverslips with cells were mounted and imaged by the investigator blinded to the cell genotype, at 20x magnification with Zeiss LSM-780 confocal microscope. Images were taken at 3-5 randomly selected areas per coverslip. For experiments involving RNF5 and RN2N signal intensity assessment, all imaging settings were kept consistent during image acquisition. For RNF5 and RN2N immunofluorescence quantification, signal intensity (mean grey value, AU) was measured in acquired images using ImageJ 2.9.0/1.53t software. For p-tau immunofluorescence quantification, p-tau-positive cells were manually counted in acquired images, blinded to cell genotype. Total cell number was quantified in the images based on nuclear Hoechst staining with the ImageJ 2.9.0/1.53t software and on average 196.66 ± 10.62 (mean ± SEM) cells were found in each image. RNF5 and RN2N signal intensity as well as p-tau positive cell number were normalised to total cell count. Measurements of both signal intensity and p-tau-positive cell quantification in each image were averaged per coverslip. Selected markers were additionally imaged at 63x and 100x magnification. Signal intensity was uniformly increased in images using ZEN Black Software (Zeiss) for presentation purposes.

### Dextran permeability assay

To assess the passive permeability of iBEC monolayers to biologically inert fluorescent tracers, cells were cultured in 0.4 μm or 3.0 μm pored Transwell inserts and fluorescein isothiocyanate (FITC)-conjugated dextran molecules of 3–5 kDa or 150 kDa (Sigma) were added at 0.5 mg/ml to the top chamber of the Transwell inserts as we previously described^41^. Following 24 h incubation with dextran, cell culture media from the bottom chamber of the Transwell system was collected (three technical replicates per Transwell) for spectrofluorometric analysis at 490 nm excitation/520 nm emission using a fluorescent plate reader (Biotek Synergy H4 or Biotek Synergy Neo2).

### Passive and FUS^+MB^-enhanced permeability of anti-tau antibodies

Monoclonal tau-specific antibodies RNF5, and RN2N, and anti-amyloid Aducanumab analogue were generated in-house as previously described^42–44,56^. Respective isotype control IgGs (IgG2b and IgG2a) were sourced from Invitrogen. For antibody permeability studies iBECs were cultured in Transwell inserts with 3.0 μm pores and exposed to selected antibody at 1 μM for 24 h as previously described by us^41^. For selected experiments, tested antibodies were conjugated with AlexaFluor-647 using Protein Labelling Kit (Invitrogen) following manufacturer’s instructions.

For FUS^+MB^-mediated antibody delivery studies, cells were exposed to 10 μl of phospholipid-shelled microbubbles with octafluoropropane gas core prepared in-house as described in^99^, together with the investigated antibodies, immediately before FUS treatment^41^. FUS was then applied at clinically relevant settings of 286 kHz center frequency, 0.3 MPa peak rarefactional pressure, 50 cycles/burst, burst period 20 ms, and a 120 s sonication time as we previously described^41,50^.

Cell culture media were collected from the bottom chamber of a Transwell system for antibody concentration assessment 24 h post antibody-or antibody and FUS^+MB^-treatment (for passive permeability and FUS^+MB^ assisted delivery assessment, respectively). For AlexaFluor-647 conjugated antibodies, the fluorescence of antibodies was measured at 633 nm excitation/665 nm emission using a plate reader (Biotek Synergy H4 or Biotek Synergy Neo2) in technical triplicate as we previously described^41^. For unconjugated antibodies, antibody concentration (ng/ml) in collected media was determined with Total Mouse IgG enzyme-linked immunoassay (ELISA, Invitrogen) and extrapolated from generated standard curves, following manufacturer instructions. For FUS^+MB^ studies, fold change in detected antibody concentration was calculated relative to its untreated (UT) control for each line at 24 h.

### RNF5, RN2N and 150 kDa dextran visualisation in *APOE* iBEC monolayers

To investigate RNF5 and RN2N localisation in *APOE* iBECs, cells were cultured in Transwell inserts with 3.0 μm pores and selected antibody conjugated to AlexaFluor647 was added at 1 μM to the top chamber of a Transwell insert for 24 h as previously described by us^41^. Following this time, the antibody was removed and cells were washed with PBS and fixed on Transwell membranes with 4% PFA for 15 min at RT. Next, cells were permeabilised with 0.3% Triton X-100 for 10 min and blocked for 1 h at RT with 2% BSA/2% GS in PBS. To enhance the signal, cells were then incubated with anti-mouse AlexaFluor647 secondary antibody (**Table S2**) diluted at 1:250 in a blocking solution for 1 h at RT in the dark. Cells were washed with PBS and Hoechst (1:5000) counterstain was performed. For dextran visualisation, iBEC were cultured in Transwell inserts and exposed to 0.5 mg/ml of FITC-conjugated 150 kDa dextran (Sigma) added to the top part of a Transwell. Following 24 h, dextran solution was removed and cells washed with PBS and fixed with 4% PFA for 15 min at RT. Cells were then washed with PBS, permeabilised with 0.3% Triton X-100 for 10 min and Hoechst (1:5000) counterstain was performed. The membranes with cells were then gently cut out of the Transwell frame with a surgical scalpel blade and mounted with Dako Mounting Medium (Agilent). Images were obtained at 20x and 63x magnification using a Zeiss LSM-780 confocal microscope. Signal intensity was uniformly increased in images using ZEN Black Software (Zeiss) for presentation purposes.

### RNA extraction, cDNA synthesis and quantitative real-time PCR (qPCR)

For *APOE* iBEC RNA collection, cells were rinsed with PBS and lysed in TRIzol^TM^ reagent (ThermoFisher Scientific) as we previously described^41,50^. Total RNA was extracted using the Direct-zol RNA Miniprep Kit (Zymo Research) according to the manufacturer’s instructions. Isolated RNA quality and quantity were measured using NanoDrop^TM^ Spectrophotometer. For quantitative real time polymerase chain reaction (qPCR) studies, 150 ng of total RNA was converted to complementary DNA (cDNA) using SensiFAST^TM^ cDNA synthesis kit following manufacturer instructions (Bioline) and qPCR performed using SensiFAST™ SYBR® Lo-ROX Kit following manufacturer instructions (Bioline). The qPCR reaction was performed in triplicate for each sample on QuantStudio^TM^ 5 Real-Time PCR system with SensiFAST™ SYBR® Lo-ROX kit-compatible cycling conditions: 2 min at 95°C followed by 40 cycles of 5 s at 95°C and 30 s at 60°C. Ct values were normalised to Ct values of *18S* endogenous control (ΔCt values). Housekeeping gene expression of *18S* was found to be consistent across cell lines. The ΔΔCt values were calculated as 2(^-ΔCt^) and presented as ΔΔCt multiplied by 10^6^. Technical triplicates were averaged per sample for statistical analysis. Utilised primer sequences are presented in **Table S3**.

### Statistical analysis

Statistical analysis was performed using GraphPad Prism version 9.4.0. Data were tested for normal distribution with Shapiro–Wilk test. For a two-group comparison with normal distribution, *F* test of equality of variances was performed and data was analysed with unpaired Student’s *t*-test (two-tailed; data with equal variances) or unpaired Welch’s *t*-test (two-tailed; data with unequal variances). Mann-Whitney *U* test (two-tailed) was used for non-normally distributed data. When comparisons between three or more groups were analysed, one-way ANOVA followed by post-hoc tests was used. *p* < 0.05 was considered statistically significant. Z-scores were calculated and values with Z-scores above or below two standard deviations (SD) of the mean were identified as outliers and excluded from analysis. Results are shown as mean ± SEM. The number of biological (*N,* hiPSC or iBEC lines) and independent (*n*) replicates used for each experiment are specified in figure legends. The number of technical replicates included in each assay is stated in the respective materials and methods sections.

## Acknowledgements

We thank QIMR Berghofer MRI Histology Facility and Microscopy Facility for their assistance. We thank Dr Satomi Okano and QIMR Berghofer MRI Statistics department for their advice on data analysis. We acknowledge Dr Carla Cuní-López for the provision of *MAPT* and *APOE* primers and Dr Gerhard Leinenga for the provision of the unlabelled Aducanumab analogue.

## Author contribution statement

**J.M.W.**: conceptualisation, methodology, investigation, formal analysis, visualisation, writing - original draft, writing - review & editing; **R.B.**: methodology, resources; **R.L.J.**: formal analysis, writing - review & editing; **J.C.S.C.**: methodology; **A.P.**: resources; **L.E.O.**: methodology; **J.G.**: methodology, resources, writing - review & editing; **R.M.N.**: conceptualisation, methodology, resources, data curation, supervision, writing - review & editing; **A.R.W.**: supervision, writing - review & editing, project administration, funding acquisition. All authors reviewed and approved the final version of the manuscript.

## SUPPLEMENTARY MATERIAL

**SUPPLEMENTAL FIGURES:**

**Supplementary Figure S1.** *APOE3*, *APOE4* and *MAPT* expression in *APOE* iBECs.

**Supplementary Figure S2.** Expression of tau detected with RNF5 antibody in *APOE* iBECs.

**Supplementary Figure S3.** Expression of tau detected with RN2N antibody in *APOE* iBECs.

**Supplementary Figure S4.** Secondary-only antibody control for RNF5 and RN2N.

**Supplementary Figure S5.** Expression of phosphorylated tau (Ser396) in *APOE* iBECs.

**Supplementary Figure S6.** Comparison of barrier integrity between iTOB0064 and TOB0064 iBECs.

**Supplementary Figure S7.** Passive permeability of anti-tau antibodies RNF5 and RN2N and anti-amyloid antibody Aducanumab in sporadic and familial AD iBECs.

**Supplementary Figure S8.** Visualisation of RNF5, RN2N and 150 kDa dextran within iBEC monolayer following 24 h.

## SUPPLEMENTAL TABLES

**Supplementary Table S1**. hiPSC lines utilised in the study.

**Supplementary Table S2.** Commercially sourced antibodies used in the study.

**Supplementary Table S3.** Primer sequences used in the study.

## Abreviations

AD: Alzheimer’s disease
APOE: apolipoprotein E
BBB: blood-brain barrier
BEC: brain endothelial cell
CSF: cerebrospinal fluid
CNS: central nervous system
ELISA: enzyme-linked immunosorbent assay
FUS: focused ultrasound
hiPSC: human induced pluripotent stem cell
iBEC: induced brain endothelial-like cell
IgG: Immunoglobulin G
MAPT: microtubule-associated protein tau gene
MB: microbubble
PSEN1: presenilin-1
p-tau: phosphorylated tau
qPCR: quantitative polymerase chain reaction
TEER: trans-endothelial electrical resistance

