## Supplemental Materials for "Increased tau expression in the *APOE4* blood-brain barrier model is associated with reduced anti-tau therapeutic antibody delivery *in vitro*"

Wasielewska *et al*.

**SUPPLEMENTAL FIGURES**

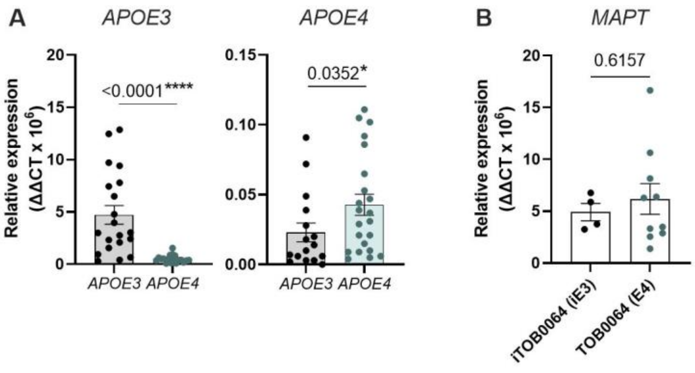


**Supplementary Figure S1. *APOE3*, *APOE4* and *MAPT* expression in *APOE* iBECs.** (**A**) Relative mRNA expression for *APOE3* and *APOE4* in utilised iBECs. $N$ = 3 biological replicates and minimum of $n$ = 5 independent replicates per line. (**B**) Relative mRNA expression for *MAPT* in iTOB0064 (*iAPOE3*) and TOB0064 (*APOE4*) iBEC. $N$ = 1 biological replicate (isogenic pair) and minimum of $n$ = 4 independent replicates per line. Results presented as $\Delta\Delta CT \times 10^6$. Data analysed with Mann-Whitney $U$ test in (A) and Student's $t$-test in (B). *$p<0.05$, ****$p<0.0001$.

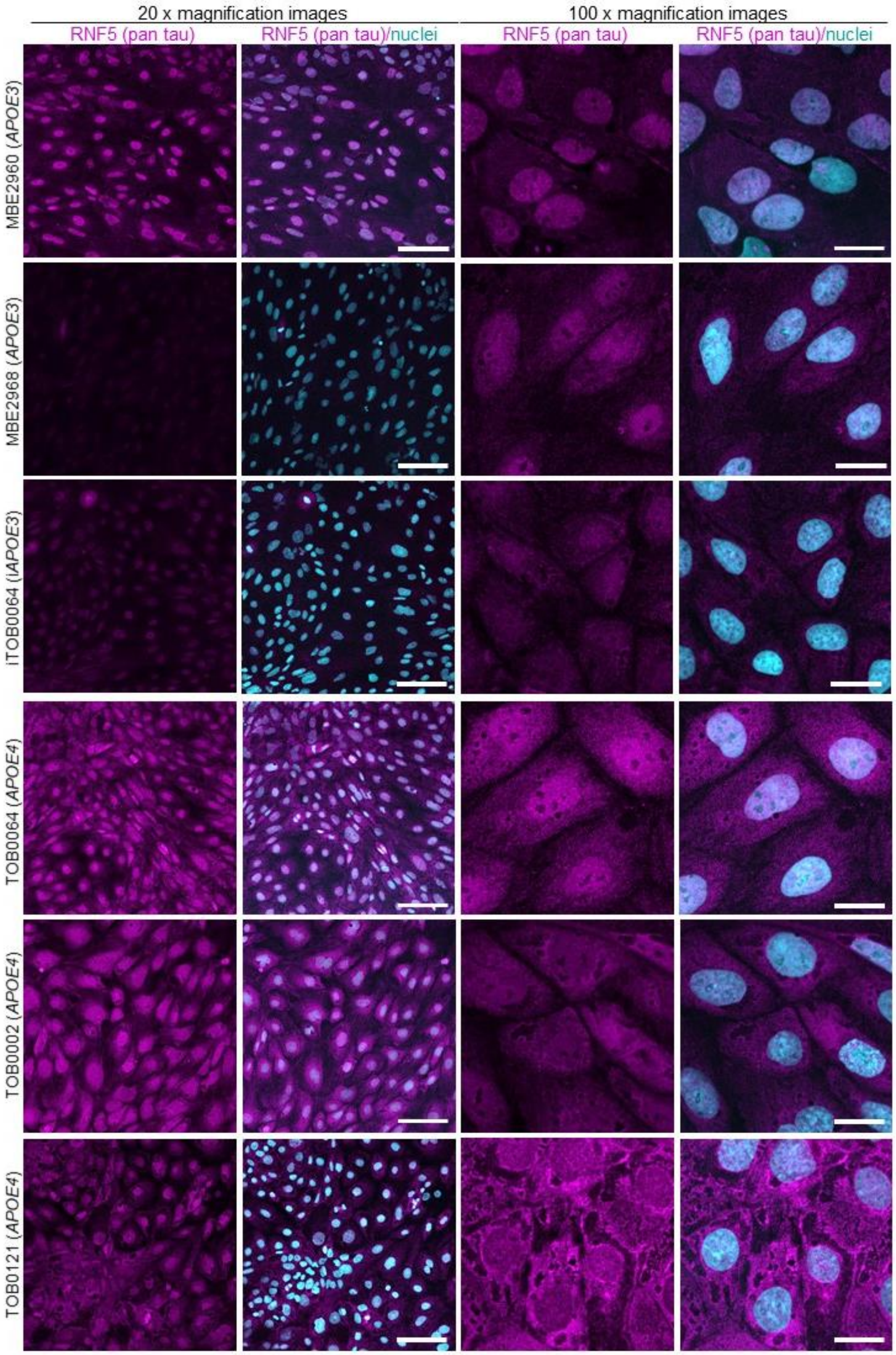


**Supplementary Figure S2. Expression of tau detected with RNF5 antibody in *APOE* iBECs.** Representative low (20x) and high (100x) magnification immunofluorescence images of tau detected by RNF5 (magenta) in *APOE3* and *APOE4* iBEC lines. Nuclei (cyan) stained with Hoechst. Scale bar represents 100 µm in 20x images, and = 20 µm in 100x images.

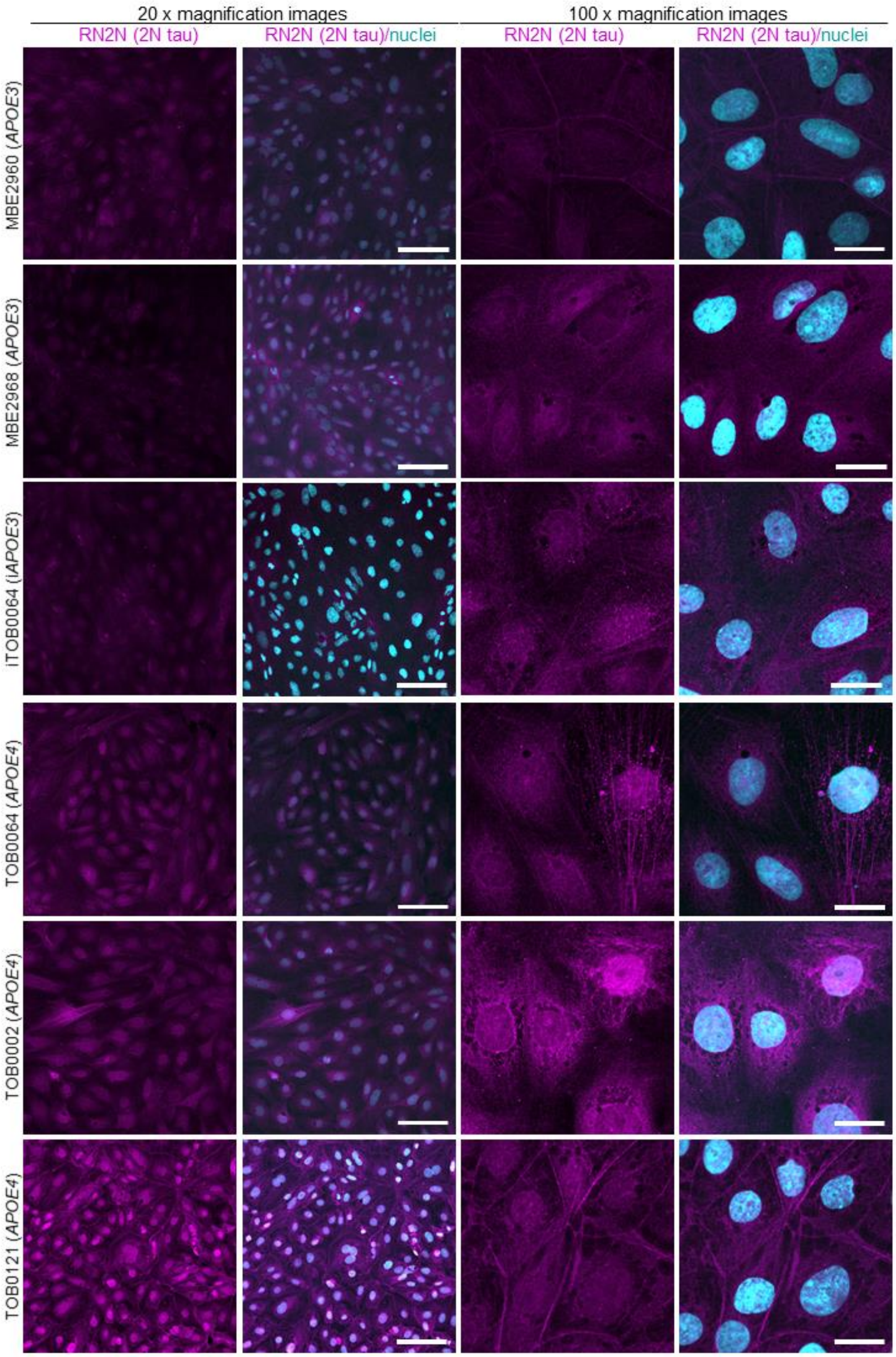


**Supplementary Figure S3. Expression of tau detected with RN2N antibody in *APOE* iBECs.** Representative low (20x) and high (100x) magnification immunofluorescence images of tau detected by RN2N (magenta) in *APOE3* and *APOE4* iBEC lines. Nuclei (cyan) stained with Hoechst. Scale bar represents 100 µm in 20x images and 20 µm in 100x images.

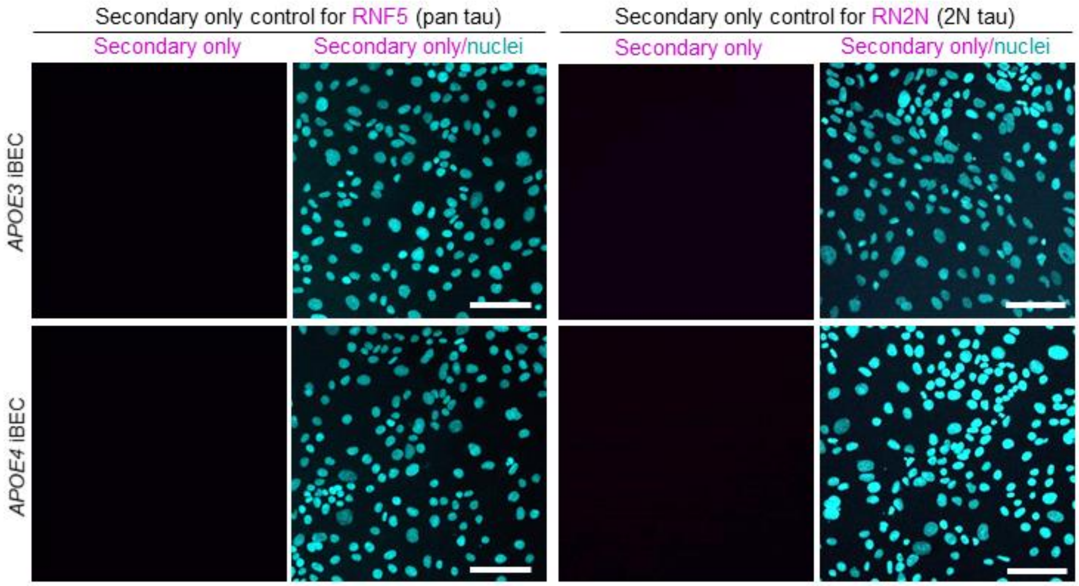


**Supplementary Figure S4. Secondary-only antibody control for RNF5 and RN2N.** Representative immunofluorescence images of secondary antibody only controls for RNF5 and RN2N (magenta) in *APOE3* and *APOE4* iBECs. Nuclei (cyan) stained with Hoechst. Scale bar = 100 µm.

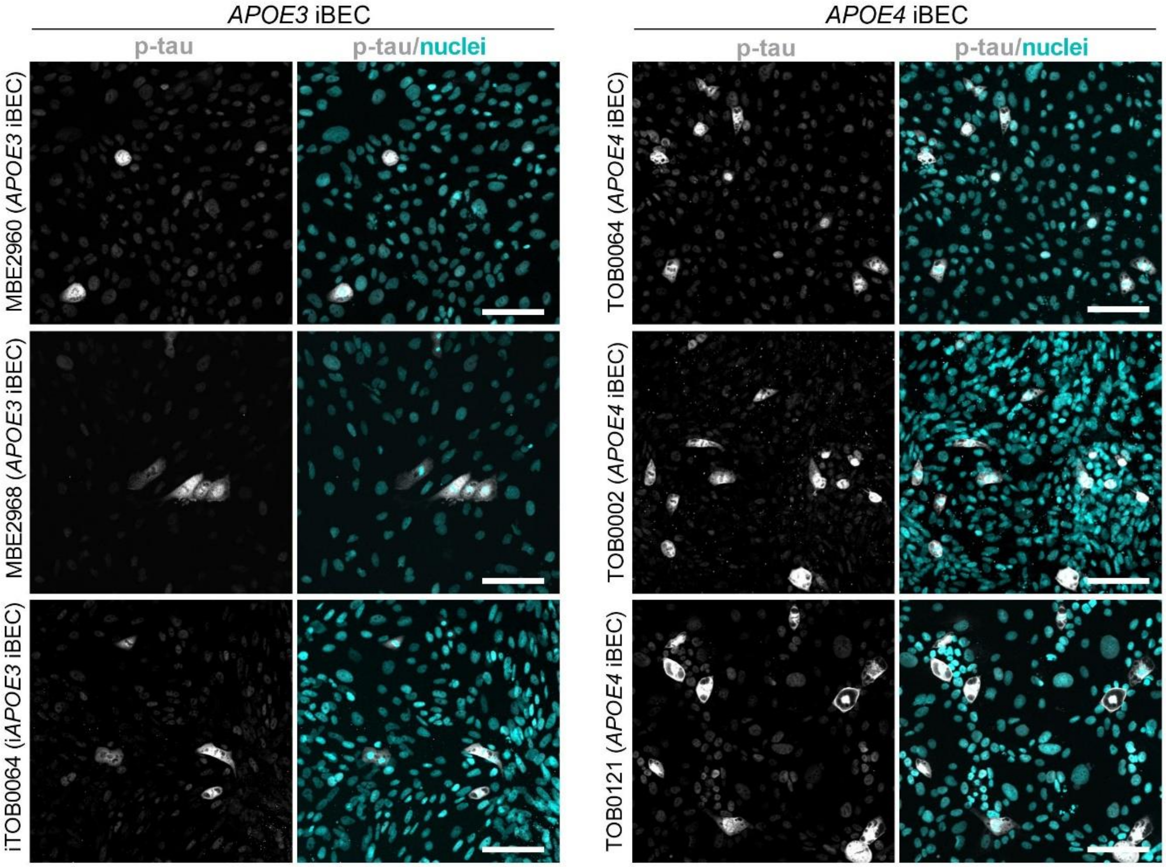

**Supplementary Figure S5. Expression of phosphorylated tau (Ser396) in *APOE* iBECs.** Representative immunofluorescence images of phosphorylated tau (p-tau, grey) with Hoechst nuclear counterstaining (cyan) in *APOE3* and *APOE4* iBEC lines. Scale bar = 100 μm.

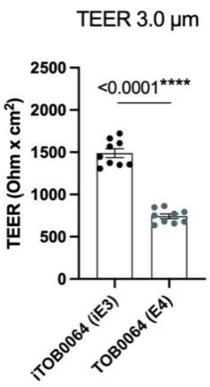


**Supplementary Figure S6. Comparison of barrier integrity between iTOB0064 and TOB0064 iBECs.** Trans-endothelial electrical resistance (TEER, shown as Ohm x $cm^2$) of iTOB0064 (*iAPOE3*) and TOB0064 (*APOE4*) iBEC, measured in Ø 3.0 µm pore Transwells. $N$ = 1 biological replicate (one isogenic pair) with $n$ = 9 independent replicates per line. Data analysed with Student's $t$-test. Error bars = SEM. ****$p$<0.0001.

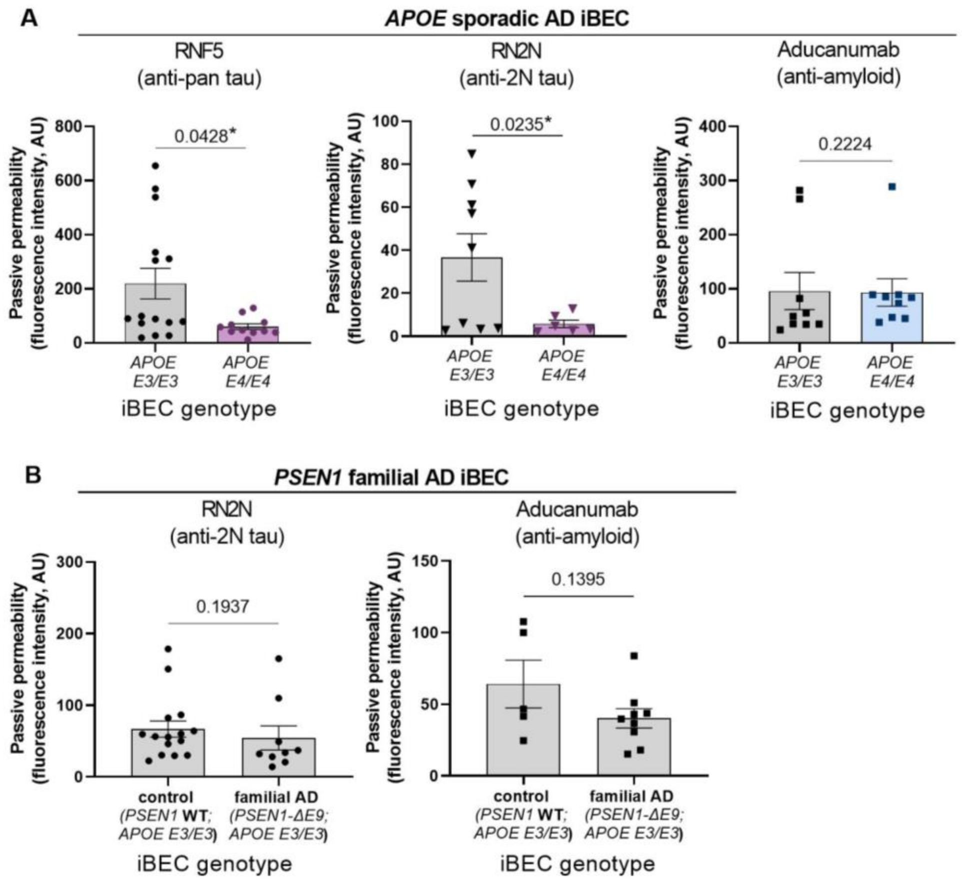


**Supplementary Figure S7. Passive permeability of anti-tau antibodies RNF5 and RN2N and anti-amyloid antibody Aducanumab in sporadic and familial AD iBECs.** (**A**) Passive permeability of AlexaFluor-647-conjugated RNF5, RN2N and Aducanumab antibodies in *APOE E3/E3* and *E4/E4* iBECs. Data presented as fluorescence intensity. RNF5: *APOE E3/E3* $N = 3$ lines, *APOE E4/E4* $N = 3$ lines and minimum of $n = 3$ independent replicates per line. RN2N: *APOE E3/E3*: $N = 3$ lines, *APOE E4/E4*: $N = 3$ lines and minimum of $n = 2$ independent replicates per line. Aducanumab: *APOE E3/E3* $N = 2$ lines, *APOE E4/E4* $N = 3$ lines and minimum of $n = 3$ independent replicates per line. (**B**) Passive permeability of AlexaFluor-647-conjugated RN2N and Aducanumab antibodies in control (*PSEN1* wildtype (WT), *APOE E3/E3*) and familial AD (*PSEN1-ΔE9* mutation, *APOE E3/E3*) iBECs. Data presented as fluorescence intensity of AlexaFluor-647-conjugated antibody measured in the bottom chamber of the Transwell at 24 h. RN2N: Control: $N = 3$ lines, familial AD: $N = 2$ lines and minimum of $n = 3$ independent replicates per line. Aducanumab: Control: $N = 3$ lines, familial AD: $N = 2$ lines and minimum of $n = 1$ independent replicate per line. Data analysed with Mann-Whitney $U$ test in (A: RNF5 and Aducanumab; B: RN2N), Student's $t$-test in (B: Aducanumab) and Welch's $t$-test in (A: RN2N). Error bars = SEM. $^{*}p<0.05$.

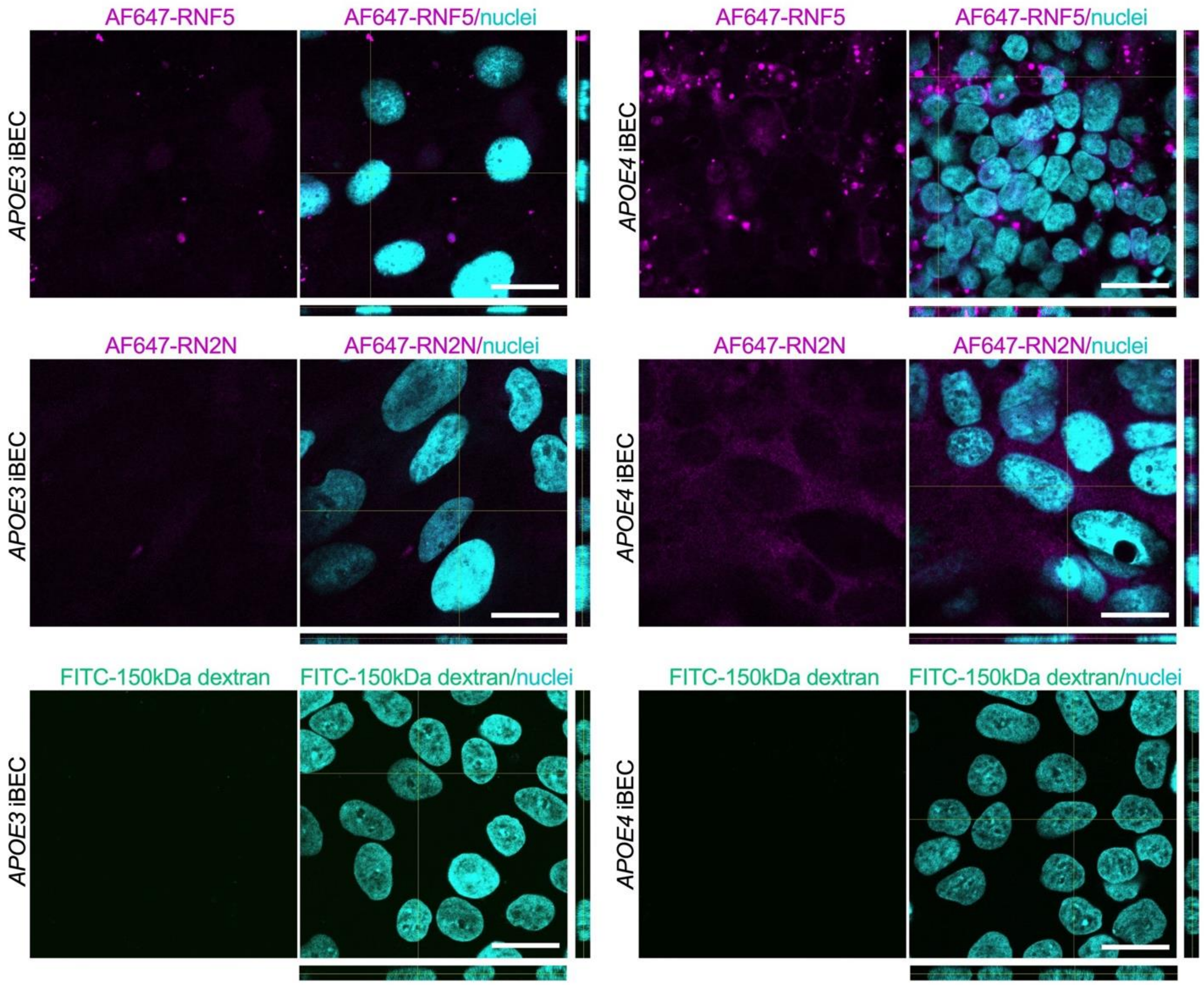


**Supplementary Figure S8. Visualisation of RNF5, RN2N and 150 kDa dextran within iBEC monolayers following 24 h.** Representative 100x magnification images showing detection of AF647-RNF5 (magenta), AF647-RN2N (magenta) and FITC-150 kDa dextran signal in iBEC monolayer formed by *APOE3* and *APOE4* cells on Transwell insert, at 24 h post-treatment with the antibody or dextran. Nuclei counterstained with Hoechst (cyan). Orthogonal views (y,z and x,z) of z-stack images are included. Scale bar = 20 μm.

### SUPPLEMENTAL TABLES

**Supplementary Table S1**. hiPSC lines utilised in the study.

<table>
<tr><th>Cell line type</th><th>Line ID</th><th>Gender</th><th>Age at biopsy</th><th>APOE genotype</th></tr>
<tr><td rowspan="6">APOE lines[1,2]:</td><td>MBE2960</td><td>Male</td><td>78</td><td>E3/E3</td></tr>
<tr><td>MBE2968</td><td>Female</td><td>65</td><td>E3/E3</td></tr>
<tr><td>iTOB0064</td><td>Male</td><td>73</td><td>E4/E4 corrected to iE3/E3 (isogenic-corrected pair to TOB0064)</td></tr>
<tr><td>TOB0064</td><td>Male</td><td>73</td><td>E4/E4</td></tr>
<tr><td>TOB0002</td><td>Male</td><td>52</td><td>E4/E4</td></tr>
<tr><td>TOB0121</td><td>Female</td><td>68</td><td>E4/E4</td></tr>
<tr><td colspan="5"></td></tr>
<tr><th>Cell line type</th><th>Line ID</th><th>Age at biopsy</th><th>PSEN1 genotype</th><th>APOE genotype</th></tr>
<tr><td rowspan="5">Familial AD lines[3–5]:</td><td>HDFa (unrelated healthy donor)</td><td>Not known</td><td>Not known</td><td>E3/E3</td></tr>
<tr><td>AD4 1.6.12.9</td><td>48</td><td>PSEN1-ΔE9 isogenic corrected</td><td>E3/E3</td></tr>
<tr><td>AD5 1.5.6.1</td><td>47</td><td>PSEN1-ΔE9 isogenic corrected</td><td>E3/E3</td></tr>
<tr><td>AD4 1.6</td><td>48</td><td>PSEN1-ΔE9</td><td>E3/E3</td></tr>
<tr><td>AD5 1.5</td><td>47</td><td>PSEN1-ΔE9</td><td>E3/E3</td></tr>
</table>

**Supplementary Table S2.** Commercially sourced antibodies used in the study.

| Antibody type | Antibody | Species | Source | Identifier |
|---|---|---|---|---|
| Primary | ZO-1 | mouse | Invitrogen | Cat#339100 |
| Primary | claudin-5 | mouse | Invitrogen | Cat#352500 |
| Primary | phosphorylated (S396)-tau | rabbit | Abcam | Cat#ab109390 |
| Primary | β-tubulin | rabbit | Cell Signaling | Cat#2146S |
| Secondary | anti-mouse AlexaFluor 488 | goat | Invitrogen | Cat#A11029 |
| Secondary | anti-mouse AlexaFluor 594 | goat | Invitrogen | Cat#A11032 |
| Secondary | anti-mouse AlexaFluor 647 | goat | Invitrogen | Cat#A32728 |
| Secondary | anti-rabbit AlexaFluor 488 | goat | Invitrogen | Cat#A11034 |

**Supplementary Table S3.** Primer sequences used in the study.

| Target gene | Forward primer sequence | Reverse primer sequence |
|---|---|---|
| *MAPT* | CTCGCATGGTCAGTAA AAGCAA | GGGTTTTTGCTGGAA TCC TGGT |
| *APOE3* | CGGACATGGAGGACGTGT | CTGGTACACTGCCAGGCG |
| *APOE4* | CGGACATGGAGGACGTGC | CGGACATGGAGGACGTGC |
| *18S* | TTCGAGGCCCTGTAATTGGA | GCAGCAACTTTAATATACGCTATTGG |
